# Uncertainty-Aware Discrete Diffusion Improves Protein Design

**DOI:** 10.1101/2025.06.30.662407

**Authors:** Sazan Mahbub, Christoph Feinauer, Caleb N. Ellington, Le Song, Eric P. Xing

**Affiliations:** GenBio AI; CMU; MBZUAI

**Author notes:** Correspondence to: Le Song < >, Eric P. Xing < >. Work done during internship at GenBio AI.

## Abstract

Protein inverse folding involves generating amino acid sequences that adopt a specified 3D structure—a key challenge in structural biology and molecular engineering. While discrete diffusion models have demonstrated strong performance, existing methods often apply uniform denoising across residues, overlooking position-specific uncertainty. We propose an uncertainty-aware discrete denoising diffusion model that employs a prior-posterior signaling mechanism to dynamically guide the denoising process. Our approach further integrates learned priors from a pretrained protein large language model and a structure encoder within a modular framework, jointly optimized through multi-objective training. Across multiple benchmarks, our method achieves substantial improvements over state-ofthe-art baselines, offering a principled framework for structure-conditioned sequence generation in proteins and beyond.

## 1. Introduction

Designing protein sequences that fold into desired three-dimensional structures is a central challenge in computational biology, with broad applications in therapeutics, synthetic biology, and molecular engineering (Dauparas et al., 2022; Wang et al., 2024; Gao et al., 2022; Sun et al., 2024; Li et al., 2014; Hsu et al., 2022). This inverse folding problem— mapping from a fixed structural scaffold to a viable amino acid sequence—presents a fundamentally multimodal and ill-posed task, where small structural variations can permit diverse valid sequences (Dauparas et al., 2022). Recent advances in deep generative modeling have shown promise in addressing this challenge, particularly through autoregressive, masked, and diffusion-based frameworks (Sun et al., 2024; Dauparas et al., 2022; Zheng et al., 2023b; Wang et al., 2024).

Among these, discrete denoising diffusion models have gained traction due to their ability to generate high-quality and realistic molecules for biomolecular design tasks (Wang et al., 2024; Ellington et al., 2024; Zou et al., 2024; Sun et al., 2024). These models simulate a corruption-recovery process over sequence space, learning to denoise structure-conditioned sequences through repeated transitions (Wang et al., 2024; Sun et al., 2024; Austin et al., 2021). However, existing formulations typically apply uniform denoising updates across all sequence positions, overlooking the fact that sequence uncertainty—often due to structural constraints— vary widely across residues and time steps. This assumption of homogeneity in uncertainty can lead to premature or unreliable updates, especially in ambiguous regions.

In this work, we introduce a novel uncertainty-aware discrete denoising diffusion model for structure-conditioned protein sequence generation. We design a prior-posterior uncertainty signaling mechanism that enables our framework to dynamically decide where and when to denoise, focusing computational effort on positions where confident updates are possible and deferring those with high residual ambiguity. This formulation enables a probabilistic and interpretable denoising trajectory that adapts to both spatial and temporal uncertainty profiles in the sequence. We also propose an approach to parameterize this formulation with a set of learnable modules that can leverage the learned prior in pretrained large language models (LLMs) and structure encoders for proteins. Furthermore, we jointly train these modules through multi-objective optimization to further enhance inverse folding performance. Our proposed approach offers a promising way to partially reduce the overhead of hyperparameter search.

Through quantitative evaluation on three widely used benchmarks, we demonstrate that our framework significantly improves upon the current state-of-the-art in structure-conditioned protein design. Beyond protein design, our uncertainty-aware diffusion framework provides a general approach for structure-conditioned discrete sequence generation and holds potential for broader applications in RNA, DNA, and other domain-specific generative modeling tasks.

## 2. Method

In this section, we derive the probabilistic model underlying our inverse folding method, starting with the simplest formulation for the denoising steps in diffusion. The problem setup is detailed in Appendix B.1.

### 2.1. Probabilistic Model for Discrete Denoising Diffusion

We seek a transition probability *P* (*x*_0_ | *x*_*T*_), which allows sampling the discrete protein sequence *x*_0_ given its noisy counterpart *x*_*T*_ . While single-step denoising has been widely studied in biological contexts (Sumi et al., 2024; Sevgen et al., 2023; Karimi et al., 2020), recent advances in iterative denoising via discrete diffusion language modeling (Ho et al., 2020; Austin et al., 2021) have introduced new directions for bio-sequence generation (Ellington et al., 2024; Sun et al., 2024; Zou et al., 2024; Wang et al., 2024). These models estimate the marginal *P* (*x*_0_ | *x*_*T*_) through intermediate variables *x*_*t*_ for *t ∈* [1, *T* −1] (Ho et al., 2020; Austin et al., 2021), with the joint probability expressed as Equation 1 and demonstrated in Figure 1a.

**Figure 1.**
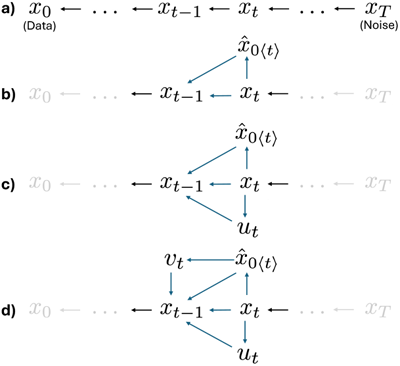
Probabilistic graphical model for the denoising process in discrete diffusion. See Section 2.1 for details.

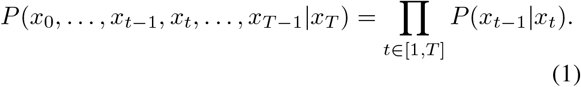

Here, *P* (*x*_*t*−1_ | *x*_*t*_) is the simplest form of a denoiser that assumes a fully Markovian reverse transition for a single reverse diffusion step from time *t* to *t* − 1, where a discrete sequence *x*_*t*_ *∈* 𝒜^*L*^ is denoised into a cleaner version *x*_*t*−1_ (here 𝒜 is a set of 20 standard amino-acids). As the noise depends on the time-step *t*, directly learning an effective transition function for *P* (*x*_*t*−1_ | *x*_*t*_) is challenging (Ho et al., 2020; Austin et al., 2021). To address this, recent works (Austin et al., 2021; Zheng et al., 2023a; Sahoo et al., 2024) introduce an intermediate variable to factorize the model. Following this strategy, we define 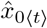 as a crude estimate of the clean data *x*_0_ given the noisy input *x*_*t*_ (Figure 1b), leading to a joint probability,

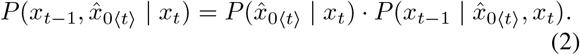

Such factorization provides us with two conditionals– 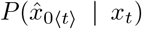 and 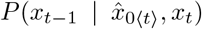. For the remaining discussion of this article, we call the former one as the denoising probability and the later as the refinement probability distribution, respectively. While it is reasonable to assume 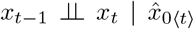 leading to 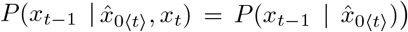, this can subsequently become a quite strong assumption about the accuracy of the samples from the denoiser 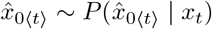. We relax this assumption in our design. Now we get the transition probability by marginalizing over 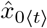,

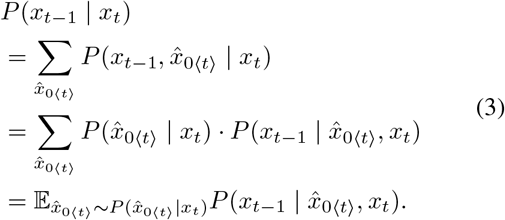

### 2.2. Uncertainty-Aware Guidance

In this section, we introduce uncertainty estimation on sequence data as a way to guide discrete diffusion. While various uncertainty estimation approaches could be integrated into our framework (Liu et al., 2020; Gawlikowski et al., 2023; Kristiadi et al., 2021; Hie et al., 2020), we follow the supervised strategy of Liu et al. (2024), motivated by its success in natural language generation. Exploring alternative techniques—including non-learnable and unsupervised methods—is left for future work.

#### Prior Uncertainty

We start by defining a random variable *u*_*t*_ *∈* {0, 1} ^*N*^, where 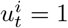 indicates the *i*-th residue is noisy, and 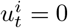 indicates it matches the native sequence (Figure 1c). Considering 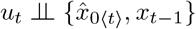, we can rewrite the joint probability in Equation 2 as,

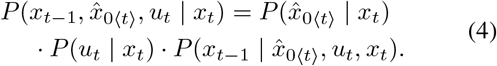

Considering *u*_*t*_ factorizes over residues,

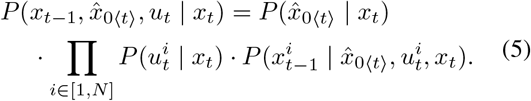

Since 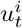 follows a Bernoulli distribution, we have 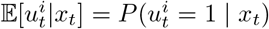, which we refer to as the *prior uncertainty estimate* and model it as a learnable function.

#### Posterior Uncertainty

We assume imperfect denoisers, sampling from which can often lead to erroneous discrete jumps. Before taking any denoising step we want to estimate how much point-wise uncertainty would change if we updated the *i*-th residue with 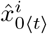. To model this, we introduce another latent variable *v*_*t*_ *∈* {0, 1}^*N*^ representing the estimated per-residue correctness of 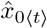, with 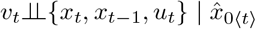 (Figure 1d). The joint probability in Equation 5 then becomes,

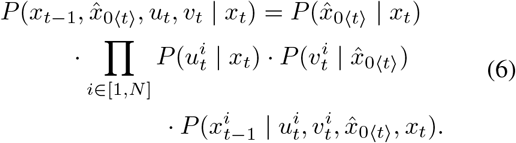

Here the expectation 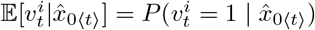 is our point-wise *posterior uncertainty* estimate.

Together, 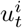 and 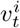 form a *prior-posterior uncertainty signal* over the *i*-th residue—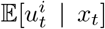 estimates existing uncertainty in 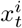, while 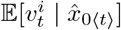 estimates uncertainty after updating with 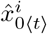. We can now marginalize over both variables to ob n the transition probability,

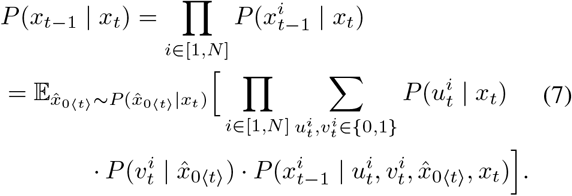

### 2.3. Conditioning on 3D Structure

Since we aim to generate a protein sequence *x*_*t*−1_ *∈ A* given its 3D conformation *ψ ∈* ℝ^*L×N×*3^, we additionally condition the transition probability *P* (*x*_*t*−1_ | *x*_*t*_) on *ψ*. This modifies the Equation as,

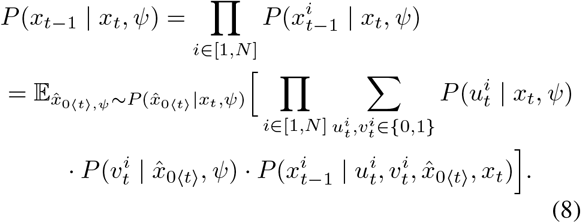

Now our prior and posterior uncertainty estimates become 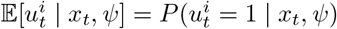 and 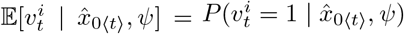, respectively.

### 2.4. Parameterization and Optimization

In this section, we describe how we parameterize the transition probability in Equation 8. We discuss our full framework as comprising five modules: (1) a structure encoder 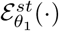, (2) a sequence encoder 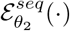, (3) a sequence de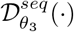, (4) an uncertainty estimator 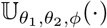, and (5) a refinement module ℛ (·). All modules except ℛ ( ·) are learnable and parameterized by *θ*_1_, *θ*_2_, *θ*_3_, and *ϕ*. We also use a single uncertainty estimator to model both prior and posterior uncertainty estimates.

#### Denoiser Parameterization

For the first three modules, we adopt AIDO.ProteinIF (Sun et al., 2024), a state-of-the-art discrete diffusion-based inverse folding method. AIDO.ProteinIF uses ProteinMPNN-CMLM (Zheng et al., 2023b) as 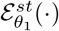, which also produces the initial sequence estimate *x*_*T*_ which is then iteratively refined, similar to other leading methods (Zheng et al., 2023b; Wang et al., 2024). 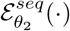 is the encoder of AIDO.Protein–a 16B parameter protein language model, while 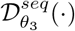 is a lightweight transformer (Vaswani, 2017) that fuses structure and sequence representations. At each time-step *t*, we obtain a crude estimate 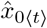 from *x*_*t*_ as,

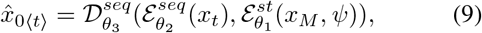

where *x*_*M*_ *∈* { <MASK>} ^*L*^ denotes a fully masked sequence with all residue labels unknown. We use *x*_*M*_ alongside the structure *ψ* as input to ProteinMPNN-CMLM, ensuring the structure encoding depends solely on *ψ*. Unlike *x*_*t*_, where each 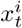 is a noisy but valid amino acid, *x*_*M*_ contains no residue information. Equation 9, representing the denoiser probability 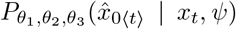, can be formulated as both deterministic or stochastic via sampling temperature (Sun et al., 2024; Wang et al., 2024). Unlike AIDO.ProteinIF, which uses time-dependent structure encoding 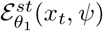, we fix it as 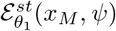, enabling caching and reducing computation. Empirically, this simplification does not affect performance.

**Figure 2.**
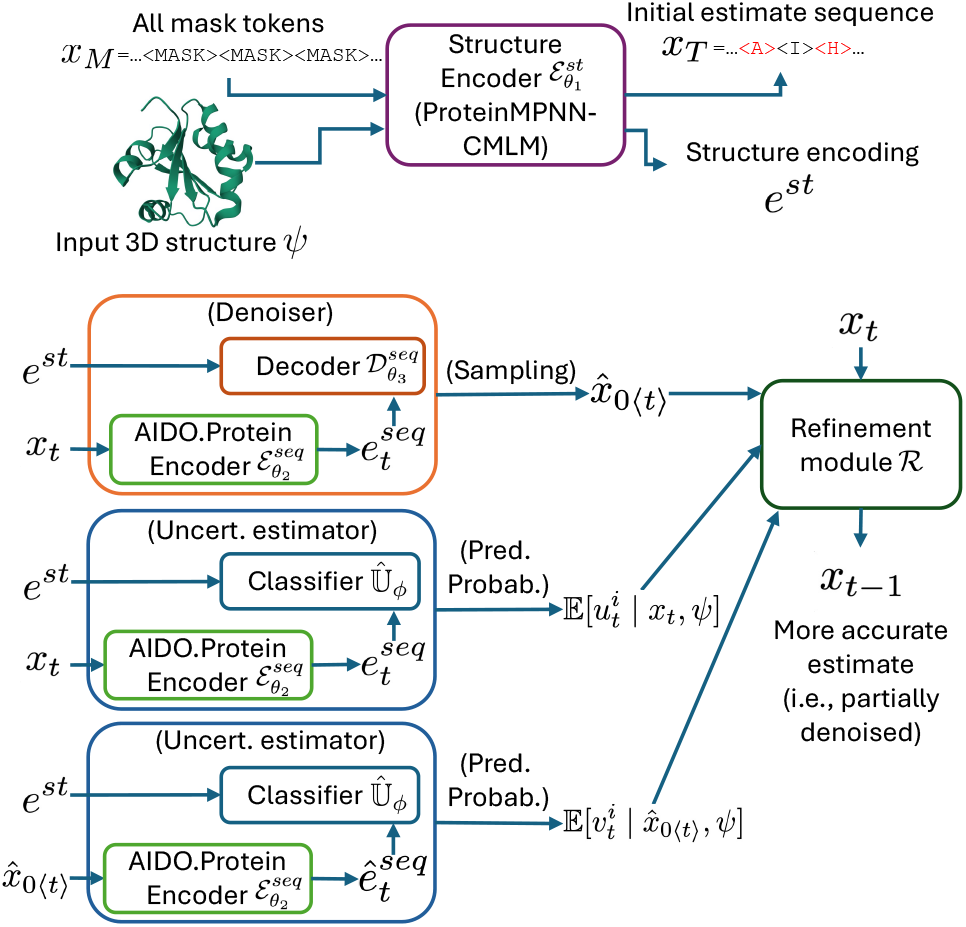
Overall architecture of our proposed method. Here, 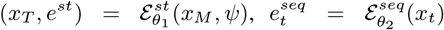, and 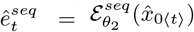. See Section 2.4 for details.

#### Uncertainty Estimator Parameterization

To leverage the learned priors from the structure and sequence encoders, we parameterize the uncertainty estimator 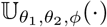 with *θ*_1_, *θ*_2_, and *ϕ*. This allows direct prediction of the prior and posterior uncertainty for residue *i* as,

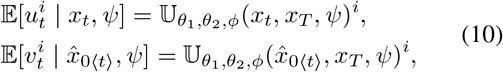

which can further be expanded as,

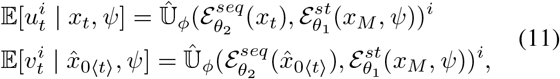

where 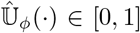 is a soft binary classifier, parameterized by *ϕ*, that uses structure and sequence encodings to predict point-wise uncertainty probabilities.

#### Refinement Module

We design the refinement module ℛ ( ·) as a simple, non-learnable function based on the estimated point-wise uncertainties,

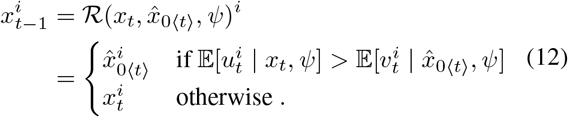

Equation 12 updates residue *i* with 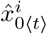 only if it reduces the estimated uncertainty. This design eliminates the need for a common hyperparameter—the number of tokens denoised per step—and makes our method less sensitive to the number of denoising steps, as uncertainty estimates serve as an implicit stopping criterion. As a result, our framework partially reduces the cost of hyperparameter tuning. In future work, we plan to explore learnable alternatives for ℛ (·).

#### Optimization

We study both individually and jointly optimized models for the denoiser (parameterized by *θ*_1_, *θ*_2_, *θ*_3_) and the uncertainty estimator (parameterized by *θ*_1_, *θ*_2_, *ϕ*). We first perform individual optimization by using the publicly available AIDO.ProteinIF model (Sun et al., 2024)^1^ and training a separate uncertainty estimator with binary classification, updating only *ϕ* while freezing *θ*_1_ and *θ*_2_. We then jointly fine-tune all parameters with a multi-objective setup combining discrete diffusion (Austin et al., 2021) and classification losses. The best performance is achieved by sampling with individually trained models and refining with jointly trained ones—consistent with prior work showing the benefit of starting from reasonable initial estimates (Sun et al., 2024; Zheng et al., 2023b; Wang et al., 2024).

## 3 Results and Discussion

We evaluate our method on standard benchmarks—CATH-4.2 (Orengo et al., 1997), TS50 (Li et al., 2014), and TS500 (Li et al., 2014)—and compare it with several state-of-the-art baselines (Appendix C.3). Results in Tables 1 and 2 report performance using perplexity (PPL) and sequence recovery (or amino acid recovery, AAR) (Zheng et al., 2023b; Wang et al., 2024); metric and dataset details are provided in Appendix C.1 and C.2.

**Table 1.**
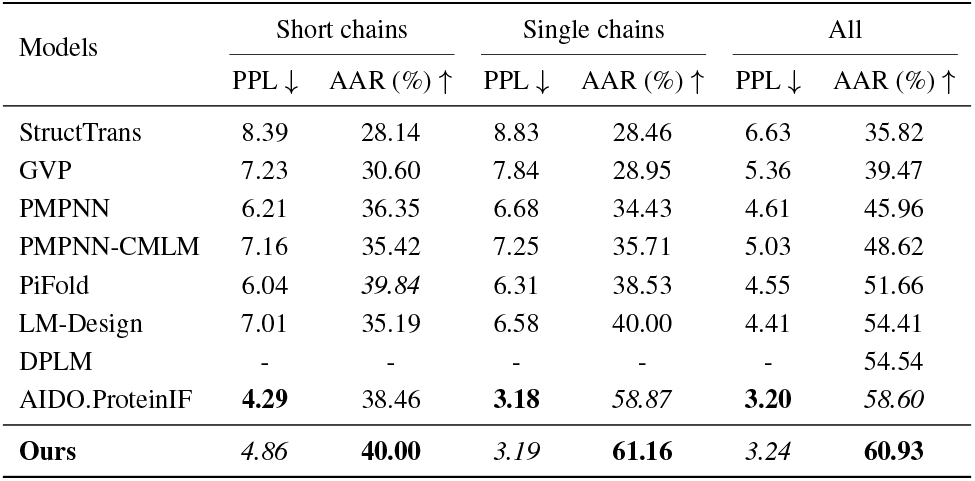
Quantitative evaluation on CATH 4.2 dataset in three different settings. Best and second best scores shown in bold and italic fonts, respectively. Here “PMPNN”= ProteinMPNN.

**Table 2.** Quantitative evaluation on TS50 and TS500 benchmark datasets. Best and second best scores shown in bold and italic fonts, respectively. Here “PMPNN”= ProteinMPNN.

| Models | TS50 |  | TS500 |  |
| --- | --- | --- | --- | --- |
|  | PPL ↓ | AAR % ↑ | PPL ↓ | AAR % ↑ |
| GVP | 4.71 | 44.14 | 4.20 | 49.14 |
| PMPNN | 3.93 | 54.43 | 3.53 | 58.08 |
| PMPNN-CMLM | 3.60 | 54.84 | 3.46 | 57.44 |
| PiFold | 3.86 | 58.72 | 3.44 | 60.42 |
| LM-Design | 3.82 | 56.92 | **2.13** | 64.50 |
| AIDO.ProteinIF | 2.93 | 66.19 | 2.68 | 69.66 |
| **Ours** | **2.86** | **68.85** | 2.59 | **71.18** |

On the CATH-4.2 dataset, our method performs consistently well across all three standard evaluation settings (Appendix C.2), achieving the highest AAR in each. It improves over AIDO.ProteinIF by 1.54% on short sequences, 2.29% on single chains, and 2.33% overall. Our method also achieved highly competitive scores in perplexity, ranking second across all CATH-4.2 settings. On the full test set, it achieves a PPL of 3.28, closely behind AIDO.ProteinIF. Investigating the divergence between PPL and AAR remains future work.

To assess generalizability, we evaluate our method on TS50 and TS500—two distinct test-only datasets (Li et al., 2014; Zheng et al., 2023b). Our model achieves state-of-the-art AAR (68.85% on TS50, 71.18% on TS500), demonstrating strong generalization across diverse proteins. It also attains the best PPL on TS50 (2.88) and the second-best on TS500 (2.62), outperforming AIDO.ProteinIF (2.68).

## 4. Conclusion

We present an uncertainty-aware discrete denoising diffusion model for structure-conditioned protein sequence generation. By employing a prior-posterior uncertainty signaling mechanism, our approach enables adaptive and interpretable denoising trajectories that account for residue- and timestep-specific ambiguity. Through modular integration with pretrained large language model and structure encoder, and joint multi-objective optimization, our method achieves significant improvements over existing baselines. These results highlight the potential of incorporating uncertainty-awareness into discrete generative frameworks for advancing sequence design across biomolecular domains.

## A. Related Work

Early breakthroughs in this space established autoregressive models as a powerful design paradigm. ProteinMPNN (Dauparas et al., 2022) exemplified this direction, achieving state-of-the-art recovery rates across a range of structural motifs and design contexts, including oligomers and binders. Its ability to generalize across diverse topologies positioned it as a widely adopted baseline. To address the computational inefficiency inherent in autoregressive sampling, PiFold (Gao et al., 2022) proposed a hybrid framework that incorporates expressive backbone encodings with an accelerated decoding scheme, delivering order-of-magnitude speedups while preserving accuracy.

Parallel to these efforts, large-scale structure-supervised training became a promising strategy. Leveraging AlphaFold2-predicted structures (John et al., 2021), Hsu et al. (2022) trained transformer architectures to directly map backbone geometries to plausible sequences, allowing the model to internalize structural priors from millions of inferred conformations. This formulation decoupled the reliance on experimental structures and enabled broader generalization.

Pretrained language models have also been adapted for inverse folding by conditioning on structure-derived features. ESM-1b and ESM-1v (Nadav et al., 2023; Meier et al., 2021) served as foundational models, later extended in Zheng et al. (2023b) for structure-guided generation. These approaches benefit from linguistic pretraining on massive protein corpora, providing rich residue-level priors that can be fine-tuned for geometry-aware decoding.

More recently, generative models based on diffusion dynamics have introduced a new formulation. Diffusion Probabilistic Language Models (DPLMs) (Wang et al., 2024) model inverse folding as a discrete denoising process, gradually refining corrupted sequences toward structure-compatible outputs. This technique introduces temporal uncertainty modeling, which facilitates a smoother posterior landscape and often yields improved diversity and stability in generation.

At the high end of model capacity, AIDO.Protein (Sun et al., 2024) utilizes a 16-billion parameter mixture-of-experts framework pretrained across multiple sequence and structure tasks. Its architecture allows conditional computation and task-specific adaptation, leading to superior performance in benchmark evaluations and improved coverage of structurally diverse regions. Given the rapidly evolving landscape, benchmarking remains a critical bottleneck. ProteinInvBench (Gao et al., 2023) was introduced to standardize evaluation across structure-conditioned generation methods. It incorporates a wide spectrum of tasks—ranging from native sequence recovery to functional design—and includes unified metrics and competitive baselines, supporting reproducibility and comparability.

## B. Preliminaries

### B.1. Problem Definition

The task of protein inverse folding involves determining a plausible amino acid sequence that would adopt a given three-dimensional (3D) structure. Let the protein conformation be denoted by *ψ* = {*ψ*^1^, *ψ*^2^, …, *ψ*^*L*^ }, where each *ψ*^*i*^ *∈* ℝ^*N×*3^ represents the spatial positions of *N* representative atoms of the *i*-th residue in 3D space, and *L* denotes the length of the protein. The objective is to predict a corresponding primary sequence *x* = [*x*^1^, *x*^2^, …, *x*^*L*^], where each *x*^*i*^ *∈* 𝒜 is an amino acid drawn from the standard set 𝒜 of canonical residues, where | 𝒜 |= 20.

The inverse folding process can be formulated as learning a function

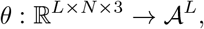

which takes the structural input and outputs a position-wise distribution over amino acids. Contemporary solutions rely heavily on neural architectures, particularly graph neural networks (GNNs) and 3D-aware models like convolutional networks adapted to irregular geometries, due to their capacity to model the intricate spatial and sequential dependencies inherent in protein structures (Gao et al., 2022; Dauparas et al., 2022; Jing et al., 2020; Hsu et al., 2022).

In many approaches, the protein structure is abstracted as a graph 𝒢 = (𝒩, *E*), where each node *n*_*i*_ *∈* 𝒩 corresponds to a residue and is annotated with geometric information (e.g., backbone coordinates), and each edge *e*_*ij*_ *∈ E* encodes relational features such as pairwise distance, angle, or biochemical interactions between residues *i* and *j* (Dauparas et al., 2022; Jing et al., 2020; Mahbub & Bayzid, 2022). This graph-based formalism allows the model to integrate local and non-local structural cues during sequence prediction.

Once trained, such models can infer a compatible sequence for a given structure either by sequentially choosing amino acids in an autoregressive fashion or by generating all residues simultaneously through a non-autoregressive mechanism, e.g., using variational autoencoders. Sampling-based techniques, including Monte Carlo simulations, and iterative refinement methods including denoising diffusion, are often used to explore diverse sequence candidates that conform to the same fold (Dauparas et al., 2022; Wang et al., 2024; Liu & Kuhlman, 2006).

## C. Experimental Setup

### C.1 Evaluation Metrics

We evaluate our model’s performance in the protein inverse folding task using two principal metrics: Perplexity (PPL) and Amino Acid Recovery (AAR).

#### Perplexity (PPL)

PPL provides a measure of the model’s predictive certainty over amino acid choices and is commonly used in sequence modeling tasks (Chen et al., 1998). In the context of protein design, lower perplexity values suggest that the model assigns high probabilities to native-like sequences, indicating that the learned distribution aligns well with that of natural proteins (Zheng et al., 2023b).

For *autoregressive* models, PPL is generally calculated as,

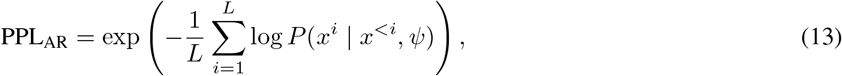

where *x*^*<i*^ denotes the sequence prefix up to position *i* − 1, and *ψ* ∈ ℝ^*L×N×*3^ represents the 3D structural context of the protein, where *L* is the number of residues in the protein and *N* is the number of representative atoms per-residue.

Since our model leverages a *non-autoregressive, iterative refinement* approach, we instead use a modified formulation where the predictions are conditioned on a noisy version of the native sequence,

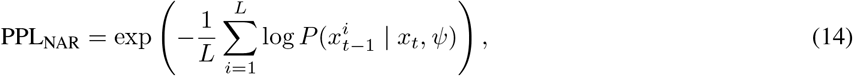

Here, *x*_*t*_ denotes a perturbed (noisy) variant of the true sequence at denoising time-step *t*, and 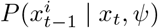 is the model’s estimated probability of observing residue 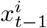 at position *i* and time-step *t* − 1 given *x*_*t*_ and *ϕ*.

#### Amino Acid Recovery (AAR)

AAR, also known as sequence recovery rate, is widely recognized as the standard metric for structure-conditioned protein design (Sun et al., 2024). It measures the typical percentage of residues in a predicted sequence that match their counterparts in the native sequence for a given protein structure. For a protein of length *L*, AAR is defined as:

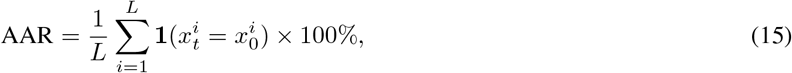

where 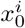 is the native amino acids at position *i* and 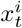 is its estimate at denoising time-step *t*, and 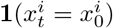 is an indicator function equal to 1 if the two residues match, and 0 otherwise. AAR is reported as the median over all the sequences in a test set (Zheng et al., 2023b; Wang et al., 2024; Dauparas et al., 2022).

### C.2. Benchmark Datasets

We conduct our experiments using three established benchmarks in the protein inverse folding literature: CATH-4.2 (Orengo et al., 1997), TS50 (Li et al., 2014), and TS500 (Li et al., 2014). Comprehensive statistics for these datasets are presented in Table 3. Among them, CATH-4.2 is a commonly adopted dataset that serves as the primary source for training, validation, and testing in numerous prior works (Gao et al., 2022; Dauparas et al., 2022; Wang et al., 2024; Sun et al., 2024). All protein sequences in this benchmark are capped at 500 amino acids in length. Within the CATH-4.2 test set, earlier studies have delineated three experimental subsets: sequences under 100 residues (termed “short sequences”), sequences that belong to a single protein chain (having only one entry in CATH 4.2, comprising roughly 92.86% of the test set), and the full test set itself. The short-sequence subset includes proteins with fewer than 100 amino acids, accounting for about 16.5% of all test samples. To evaluate generalization beyond the training domain, we additionally assess performance on TS50 and TS500. TS50 contains only 50 proteins, with the longest sequence spanning 173 residues. In contrast, TS500 introduces significant sequence length diversity, ranging from just 43 to as long as 1,636 residues. Following prior protocols (Zheng et al., 2023b; Gao et al., 2022), we use TS50 and TS500 exclusively for evaluation, relying on models trained solely on the training portion of CATH-4.2.

**Table 3.** Overview of dataset composition and sequence length distribution across CATH-4.2, TS50, and TS500. Columns: “Sample count” = number of sequences, “Residue count” = number of total amino acids, “Avg.” = average sequence length, “Med.” = median sequence length, “S.D.” = standard deviation of sequence lengths.

| Dataset | Split | Sample count | Residue count | Avg. | Med. | S.D. |
| --- | --- | --- | --- | --- | --- | --- |
| CATH-4.2 | Train | 18,024 | 3,941,775 | 218.7 | 204.0 | 109.93 |
|  | Validation | 608 | 105,926 | 174.2 | 146.0 | 92.44 |
|  | Test | 1,120 | 181,693 | 162.2 | 138.0 | 82.22 |
|  | All | 19,752 | 4,229,394 | 214.1 | 196.0 | 109.06 |
| TS50 | Test | 50 | 6,861 | 137.2 | 145.0 | 25.96 |
| TS500 | Test | 500 | 130,960 | 261.9 | 225.0 | 167.30 |

### C.3. Baselines for Comparison

To benchmark our method, we compare against eight state-of-the-art approaches that exemplify diverse modeling paradigms for structure-conditioned protein sequence generation: AIDO.Protein (Sun et al., 2024), ProteinMPNN (Dauparas et al., 2022), its non-autoregressive counterpart ProteinMPNN-CMLM (Zheng et al., 2023b), LM-Design (Zheng et al., 2023b), DPLM (Wang et al., 2024), PiFold (Gao et al., 2022), GVP (Jing et al., 2020), StructTrans (Ingraham et al., 2019).

AIDO.Protein (Sun et al., 2024) leverages large-scale pretraining—spanning 16 billion parameters—and adapts to the inverse folding task through a discrete diffusion modeling objective that is explicitly conditioned on structural input. ProteinMPNN (Dauparas et al., 2022), an autoregressive model, generates amino acid sequences sequentially based on structural input, whereas ProteinMPNN-CMLM (Zheng et al., 2023b) replaces the autoregressive decoding with a masked prediction strategy via the conditional masked language modeling (CMLM) objective (Ghazvininejad et al., 2019), yielding improved sequence recovery. Built similarly on the CMLM framework, LM-Design (Zheng et al., 2023b) augments non-autoregressive decoding with protein language model pretraining to further boost performance on inverse folding. DPLM (Wang et al., 2024) takes a different direction by introducing discrete diffusion objectives, enabling more flexible sequence modeling through iterative refinement. PiFold (Gao et al., 2022) combines expressive structural features with efficient autoregressive decoding, offering both competitive accuracy and inference speed. GVP (Jing et al., 2020) extends classical neural architectures by incorporating geometric vector perceptrons, enabling operations directly on 3D Euclidean features. StructTrans (Ingraham et al., 2019) adopts a conditional generation framework over protein graphs to synthesize sequences compatible with given backbone structures.

## Footnotes

1 https://huggingface.co/genbio-ai/AIDO.ProteinIF-16B

